# CloudASM: an ultra-efficient cloud-based pipeline for mapping allele-specific DNA methylation

**DOI:** 10.1101/2020.01.28.887430

**Authors:** Emmanuel LP Dumont, Benjamin Tycko, Catherine Do

**Affiliations:** Hackensack-Meridian Health Center for Discovery and Innovation, Nutley, NJ 07110, USA; Hackensack-Meridian Health School of Medicine at Seton Hall University, Nutley, NJ 07110, USA; Lombardi Comprehensive Cancer Center, Georgetown University, Washington, DC 20007

## Abstract

**Summary:** Methods for quantifying the imbalance in CpG methylation between alleles genome-wide have been described but their algorithmic time complexity is quadratic and their practical use requires painstaking attention to infrastructure choice, implementation, and execution. To solve this problem, we developed CloudASM, a scalable, ultra-efficient, turn-key, portable pipeline on Google Cloud Computing (GCP) that uses a novel pipeline manager and GCP’s serverless enterprise data warehouse.

**Availability and Implementation:** CloudASM is freely available in the GitHub repository https://github.com/TyckoLab/CloudASM and a sample dataset and its results are also freely available at https://console.cloud.google.com/storage/browser/cloudasm.

**Supplementary information:** None.

## 1 Introduction

The importance of allele-specific methylation (ASM) at CpG sites was recently illustrated by linking haplotype-dependent ASM to genetic variants (single-nucleotide polymorphisms; SNPs) that disrupt transcription factor binding sites (Do *et al.*, 2016, 2017, 2019; Onuchic *et al.*, 2018). Correctly mapping ASM in whole methylomes (e.g. data from whole-genome bisulfite sequencing) requires the joint analysis of three gigabyte-large datasets, namely, aligned reads, variant calling on these reads, and CpG methylation calling on these reads. Therefore, implementing such a pipeline at scale becomes a demanding bioinformatics endeavor involving irreversible decisions on infrastructure, implementation, and execution. Current approaches to call ASM (Onuchic *et al.*, 2018; Orjuela *et al.*, 2019; Do *et al.*, 2016, 2019) are similar in that they create intertwined loops on the three datasets of aligned reads, variant calling, and CpG methylation calling, which lead to several operations with a quadratic algorithmic time complexity. For instance, finding the genotyping of each variant in each read has a quadratic time complexity of *O(NxM)* where *N* is the number of reads and *M* the number of variants. Therefore, such operations require an access to a large number of CPUs if these loops are parallelized over smaller datasets, preventing researchers from systematically mapping allele-specific methylation in bisulfite-converted genomes.

## 2 Presentation of CloudASM

### 2.1 Overview

To address the issues described in the introduction, we designed CloudASM, a cloud-based, scalable, fully-integrated and highly-efficient pipeline to call ASM from fastq files resulting from bisulfite sequencing. CloudASM enables anyone to map ASM on any number of samples in the same amount of time and from any machine, provided they have an account with GCP with the appropriate authorizations and quotas (see our Github repository for more details). CloudASM uses Google’s pipeline manager “dsub”, which handles workloads and job submission similarly to slurm or Grid Engine (Kohlhoff, 2019). Also, like recent pipeline managers (Lee *et al.*, 2019), dsub accepts reproducible and portable pipeline standards (e.g. Docker) and cloud configuration is automatically handled with a serverless scheduling approach.

### 2.2 Optimization of ASM-specific steps

As described in the introduction, steps specific to mapping ASM exhibit a quadratic complexity, an expensive endeavor in terms of CPU-hours. To break with this approach, we used GCP’s “BigQuery” module for ASM-specific steps. Created internally by Google to manipulate its own large datasets through a severless and scalable data warehouse, and made commercially-available in 2010, BigQuery is ideally suited to elasticaly manipulate and analyze large genomic datasets. Specifically, we designed ASM-specific steps using BigQuery’s join function. Although BigQuery’s join algorithm is not publicly available, peer-reviewed algorithms for join functions were shown to have a linear or quasilinear time complexity (Chen and Zhong, 2014), representing a significant improvement over the quadratic time complexity found in current ASM scripts. We further minimized the computation time by selectively using “Broadcast” and “Hash” join queries (Lakshmanan and Tigani, 2019). After implementing ASM-specific on BigQuery, we measured a 25x improvement in CPU-hours compared to the internal pipeline that we have used for previous publications (Do *et al.*, 2017, 2019, 2016) and that uses the algorithmic approach of intertwined loops on the aligned reads, variant calling, and CpG methylation calling. See section 3.4 for details regarding the comparison.

### 2.3 Adjustable parameters

CloudASM can be customized using 10 different adjustable parameters, covering alignment, variant call, and the stringency of ASM regions. The user can choose which reference genome will be used for aligning the fastq files (hg19 or GrCh38) and the script will automatically download the selected reference genome, its most recent associated variant database, and convert the reference genome into a bisulfite genome. The user can also choose which database of variants should be used to remove CpG sites that may overlap with variants (unfiltered or filtered variants derived from the variant call on the aligned reads, or the database of common variants derived from the most updated variant database associated with the selected reference genome, with the option of selecting the frequency of common variants - 0.05 by default). When performing the task of identifying the reads that include a biallelic variant, the user can select the minimum quality score recalibrated by the variant call required to identify the variant (zero by default). To map ASM regions, CloudASM computes ASM at the CpG level and across reads overlapping the variant, ensuring that ASM regions identified by CloudASM are not artifacts of misalignment and methylation/variant call errors.

First, the user can define the effect size of the difference between fractional methylation levels across reads between the two alleles (20% by default), the minimum coverage per CpG across reads per allele (5x by default), the minimum number of CpG sites “near” a variant to test if there is ASM (3 per default), and the minimum number of CpG sites with a significant methylation difference and in the same direction between the two alleles (3 per default). The user can also select the minimum number of consecutive CpG sites with a significant methylation difference (2 by default), which we found to be the most stringent parameter when calling ASM. Also, when calculating the effect size of an ASM region and its p-value using a Wilcoxon test, the user can select the p-value threshold for significance (0.05 by default) or the Benjamini/Hochberg threshold (0.05 by default) used when correcting the p-value for multiple testing. Finally, when calculating ASM at the CpG level and its significance using a Fisher test, the user can select the p-value for its significance (0.05 by default).

**Figure 1.**
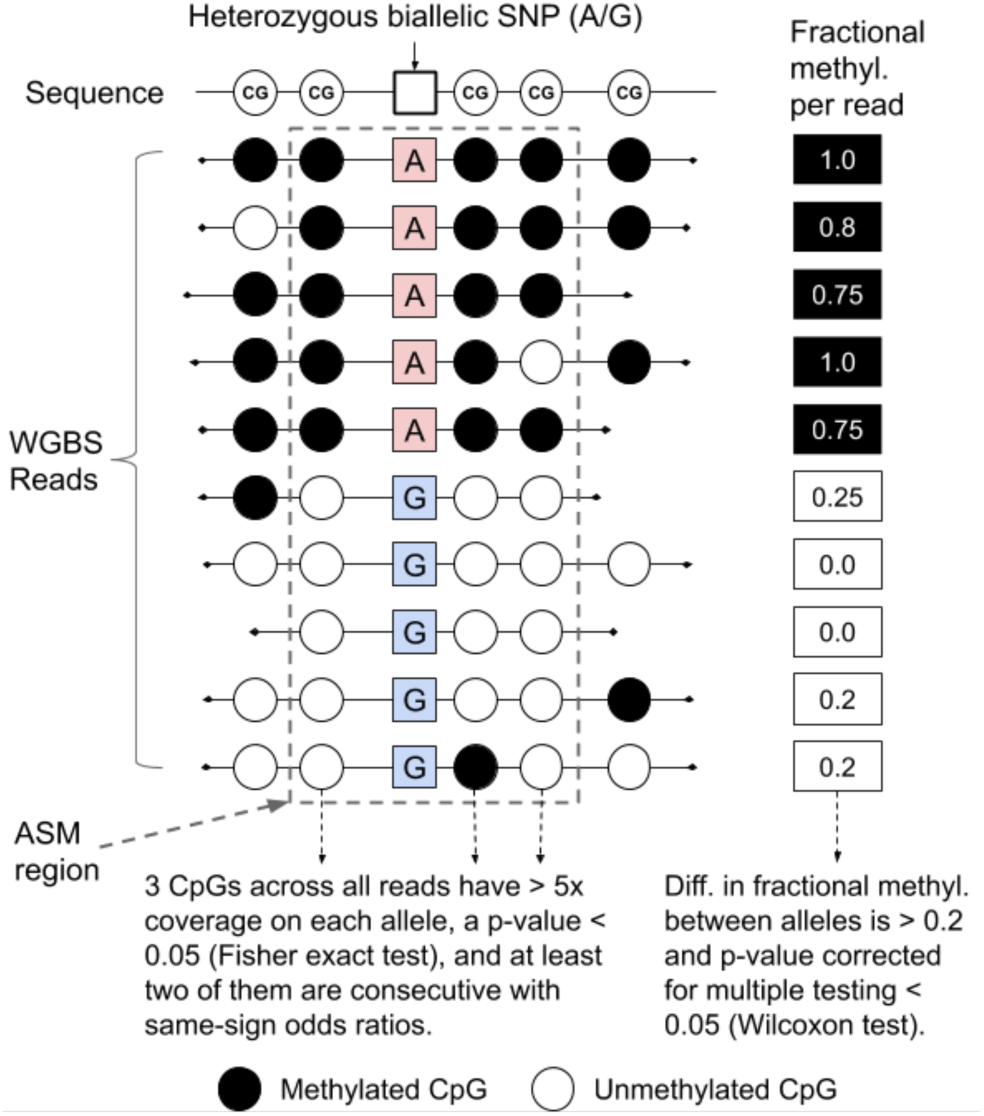
Definition of ASM in CloudASM. All parameters are customizable by the user. Cloud ASM uses a 2-dimension (by CpGs and across CpGs) and a 2-step statistical approach to define ASM.

## 3 Results

### 3.1 How to use CloudASM

Our pipeline can be run from any machine that is equipped with a command line terminal and Python (nearly all PC, Mac, and Linux machines). To employ our pipeline, a user must install Google’s pipeline manager dsub and have access to BigQuery, Compute Engine, and Cloud Storage in a GCP project. The user does not need to install any bioinformatics package because CloudASM uses our publicly-available Docker images which come with all the packages required by CloudASM. If the user is new to GCP, we provide additional information on our Github repository.

### 3.2 Execution example

We prepared a small dataset (∼24MB of zipped fastq files) for anyone to rapidly test CloudASM without incurring large costs, prior to applying it to whole methylomes. The dataset was prepared by recombining bisulfite-converted reads overlapping the positions from 9,000,000 to 10,000,000 on chromosome 1, using the module *bamtofastq* within the package BEDTools (Quinlan and Hall, 2010) on the lymphoblastoid cell line GM12878 (identifier: ENCSR890UQO) made available by the ENCODE consortium (Davis *et al.*, 2018). The zipped fastq files are freely accessible on our GCP’s storage space at https://console.cloud.google.com/storage/browser/cloudasm. Using this dataset, CloudASM assessed 456 SNPs and found 13 ASM regions. All the data generated by CloudASM is stored in the same location in our GCP storage space.

Finally, we ran both CloudASM and a traditional cloud-powered cluster using a cluster manager (Elasticluster) and job scheduler (Sun Grid Engine, SGE) on all fastq files for GM12878 and measured the number of CPU-hours in both cases. Both algorithms evaluated 1.5 billion reads, 1.1 billion CpG methylation status, 1.48 million variants and found 8,045 ASM regions. We measured a 25x improvement in the number of CPU-hours of CloudASM over the cloud-powered cluster we used in (Do *et al.*, 2019).

**Table 1.** Comparison between CloudASM and the pipeline used in (Do *et al.*, 2019) to call ASM on ENCODE’s lymphoblastoid cell line GM12878 (identifier: ENCSR890UQO on the ENCODE website).

| ASM Pipeline | CloudASM | SGE-powered pipeline |
| --- | --- | --- |
| Number of CPU-hours | 77 | 1,944 |

### 3.4 Scalability of CloudASM

Because CloudASM uses Google’s pipeline manager dsub, CloudASM does not rely on a cluster (*i.e.* with a manager node and a set of compute nodes communicating through a NFS server) and, therefore, its scalability is not limited by common issues found in clusters, such as NFS I/O issues and cluster management. In clusters (on-site or cloud-powered), compute nodes share a common drive where they perform operations and eventually, they collectively reach the disk’s maximum writing speed. We noticed this issue both using Elasticluster/SGE and HTCondor on GCP. Finally, clusters need to be created, maintained, and their nodes need to be disposed of when they are no longer used. This servicing to cluster is both time and resource consuming. CloudASM, through dsub, automatically creates, maintains, and kills virtual machines with their own disks, eliminating scalability issues encountered with cluster management. To be scalable over any number of samples, the user will have to make sure that her project has been authorized by GCP to request the resources required by CloudASM (See our Github repository for more info). We have used CloudASM to process 10 whole methylomes in parallel and the duration of their analysis was not noticeably different from the time required to analyze a single whole methylome. However, the duration to process 10 whole methylomes on a cloud-powered SGE cluster increases approximately by 50% because of the issues described above, therefore increasing the number of CPU-hours by 50% as well.

### 3.5 The use of preemptible machines

CloudASM has a few steps relying on preemptible virtual instances (e.g. trimming of the reads). These machines are about 80% cheaper than standard virtual machines but they live for at most 24 hours and are subject to random interruptions. On average, we observe a 15% failure rate in a given batch relying on these machines. If the user does not wish to manage these failed jobs, she should delete every row containing “--preemptible” in the master.sh script (The dsub pipeline manager does not let users choose if their virtual instances are preemptible or not using a variable so we could not make it an adjustable parameter). If the user wishes to make CloudASM more affordable by using preemptible instances, we provide the code we have been using to re-run the jobs that failed in our Github repository.

## 4 Conclusion

CloudASM is a cloud-based, highly-efficient pipeline to call ASM. We believe that the unprecedented efficiency of our pipeline will give researchers easy access to a useful regulatory epigenome mark and that its flexibility will be useful in exploring other allele-specific epigenomes or transcriptomes.

## Acknowledgements

We are grateful to Peter Kaplan for his feedback on the manuscript. We also thank Adil Alaoui and Subha Madhavan at Georgetown University for setting up a Google Cloud Computing account on our behalf. Finally, we thank Jesus Gomez and Keith Binder at Google for their advice.

